# GenMap: Fast and Exact Computation of Genome Mappability

**DOI:** 10.1101/611160

**Authors:** Christopher Pockrandt, Mai Alzamel, Costas S. Iliopoulos, Knut Reinert

## Abstract

We present a fast and exact algorithm to compute the (*k, e*)-mappability. Its inverse, the (*k, e*)-frequency counts the number of occurrences of each *k*-mer with up to *e* errors in a sequence. The algorithm we present is a magnitude faster than the algorithm in the widely used GEM suite while not relying on heuristics, and can even compute the mappability for short *k*-mers on highly repetitive plant genomes. We also show that mappability can be computed on multiple sequences to identify marker genes illustrated by the example of E. coli strains. GenMap allows exporting the mappability information into different formats such as raw output, wig and bed files. The application and its C++ source code is available on https://github.com/cpockrandt/genmap.

## 1. Motivation

Analyzing data derived from massively parallel sequencing experiments often depends on the process of genome assembly via re-sequencing; namely, assembly with the help of a reference sequence. In this process, a large number of reads derived from a DNA donor during these experiments must be mapped back to a reference sequence, comprising a few gigabases to establish the section of the genome from which each read originates. An extensive number of short-read alignment techniques and tools have been introduced to address this challenge emphasizing different aspects of the process [1]. In turn, given a set of reads of some fixed length *k* the process of re-sequencing depends heavily on how mappable a genome is. Thus, for every substring of length *k* in the sequence we want to count how many times this substring appears in the sequence while allowing for a small number *e* of errors. A great variance in genome mappability between species and gene classes was revealed in [2].

The concept of mappability for sequence analysis was introduced by Koehler et al. [3], taken up again and later formalized by Derrien et al. [2] (see also [4]). They also implemented an algorithm based on a heuristic to compute the mappability as part of the GEM tools.

### Definition ((*k, e*)-mappability and (*k, e*)-frequency)

Given a string *T* of length *n*, the (*k, e*)-frequency counts occurrences of every single *k*-mer in *T* with up to *e* errors. We denote the *k*-mer starting at position *i* in *T* as *T*_*i*_. The values are stored in a frequency vector *F* of length *n - k* + 1 such that

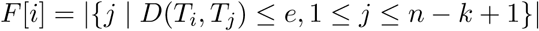

where *D*(*T*_*i*_, *T*_*j*_) denotes the distance of two *k*-mers given a metric such as Hamming or Edit distance. Its elementwise multiplicative inverse is called the (*k, e*)-mappability and stored in a mappability vector *M* with *M* [*i*] = 1*/F* [*i*] for 1 *≤ i ≤ n - k* + 1.

Intuitively speaking a mappability value of 1 represents a unique *k*-mer, a mappability value close to 0 indicates a *k*-mer occurring in repetitive regions. Figure 1 gives an example for the (4, 0) and (4, 1)-frequency of a given text.

**Figure 1:**
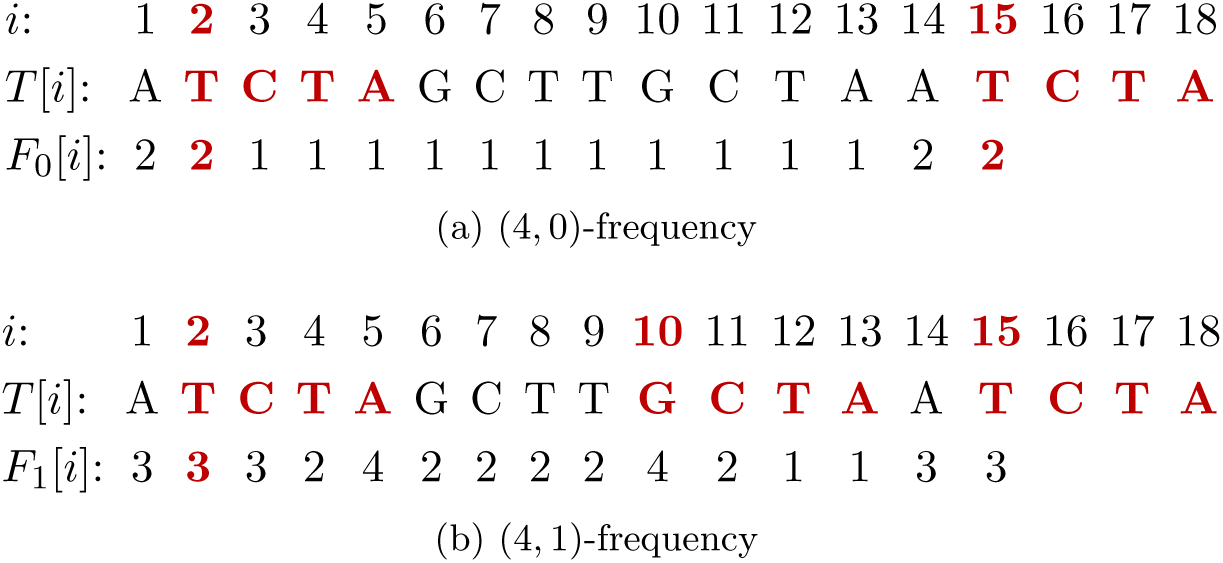
(*k, e*)-frequency vectors *F*_*e*_ for *k* = 4 and *e* ∈ {0, 1} on the same sequence. A frequency of 1 indicates that the *k*-mer starting at that position in the text is unique in the entire sequence without errors respectively with up to one mismatch.

Since for some applications an exact computation of mappability is favorable, we propose a new algorithm that is not only faster than previous ones, but also exact, i.e., without approximating mappability values. Mappability can not only be used straightforward to retrieve information on the repetitiveness of the underlying data. In this paper we will also illustrate that it can be used to find marker sequences that allow distinguishing similar strains of the same species, as well as separate strains by groups sharing common *k*-mers.

## 2. Algorithm

Before we present our algorithm we give an overview on the approach of Derrien et al. to compute the (*k, e*)-mappability. For reasons of clarity we consider computing its inverse, the (*k, e*)-frequency and neglect searching the reverse strand throughout this paper. To consider the reverse strand, each *k*-mer has simply to be searched by its reverse complement leading to a doubling of the running time. Furthermore, we consider Hamming distance, if not stated otherwise. It can be applied to other distance metrics such as Edit distance as well.

To achieve a feasible running time Derrien et al. implemented a heuristic to approximate some of the frequency values. First, they initialize the frequency vector with 0s and perform a linear scan over the text (see algorithm 1). Then each *k*-mer *T*_*i*_ is searched with *e* errors in an FM index and the number of occurrences is stored in *F* [*i*]. If the count value exceeds some user-defined threshold parameter *t*, the locations of these occurrences are located. Let *j* be such a location. Since *T*_*i*_ has a high frequency, i.e., *F* [*i*] *> t* and *D*(*T*_*i*_, *T*_*j*_) *≤ e*, it is likely that *T*_*i*_ and *T*_*j*_ share common approximate matches. Hence, *F* [*j*] is assigned the frequency value *F* [*i*]. To speed up the computation, *k*-mers that already have frequency values assigned due to this approximation are skipped during the scan over the text. If a position *j* is located multiple times as an approximate match of a repetitive *k*-mer, *F* [*j*] is assigned the maximum frequency of all these *k*-mers to avoid underestimating the frequency value *F* [*j*].

### Algorithm 1 Inexact algorithm to compute the (*k, e*)-frequency by Derrien et al

1: **procedure** INEXACT FREQUENCY(*T, k, e, t*)

2: *F* [1..*|T | - k* + 1] *←* {0}

3: **for** *i* = 1, *…, |F |* **do**

4: **if** *F* [*i*] = 0 **then**

5: *F* [*i*] *← |𝒫|*

6: 𝒫 *←* **approximate matches with** *e* **errors**

7: **if** *|𝒫| > t* **then**

8: **for** *j ∈ 𝒫* **do**

9: *F* [*j*] *←* max(*F* [*j*], *|𝒫|*)

10: **return** *F*

Their experiments on chromosome 19 of the human genome with *t* = 7 show that almost 90 % of the 50-mers with a frequency of 3 are correct, for 50-mers with frequency values between 8 and 12 only 75 % are correct (similar errors for C. elegans with *t* = 6). This can be led back to an overestimation of rather unique k-mers.

We now present a fast and exact algorithm to compute the (*k, e*)-frequency. Similar to the algorithm implemented in GEM we scan over the text *T* while searching and counting the occurrences of each *k*-mer *T*_*i*_ for 1 *≤ i ≤ n -k* + 1 with up to *e* errors in an index on *T*. In contrast to GEM we improve the running time by reducing redundant searches with three major improvements which we introduce in the following.

### 2.1 Adjacent k-mers

Adjacent *k*-mers in *T* are highly similar, since they have a large overlap. Hence, we do not search every *k*-mer separately. Consider the adjacent *k*-mers *T*_*j*_, *T*_*j*+1_, *…, T*_*j*+*s-*1_ for some integer *s ≤ k - e* + 1 which all share the common sequence *T* [*j* + *s -* 1..*j* + *k -* 1] of length at least *e*. Since we already need to allow for up to *e* errors in their common sequence when searching each *k*-mer, this infix should only be searched once. Thus, we start with searching this infix and extend it afterwards to retrieve the occurrences for each *k*-mer separately.

A search in an index is performed character by character. Unidirectional indices only support extending characters into one direction, either to the left or to the right, while bidirectional indices support searching into both directions in any arbitrary order [5]. To perform approximate string matching, backtracking is performed to search for every possible approximate match within the given error bound. First, backtracking with up to *e* errors is performed to search the infix and afterwards it is extended to search each *k*-mer separately using backtracking allowing for the remaining number of errors not spent in the search of the infix. Since the extension is performed into both directions, a bidirectional index is required. Figure 2a illustrates this approach.

**Figure 2:**
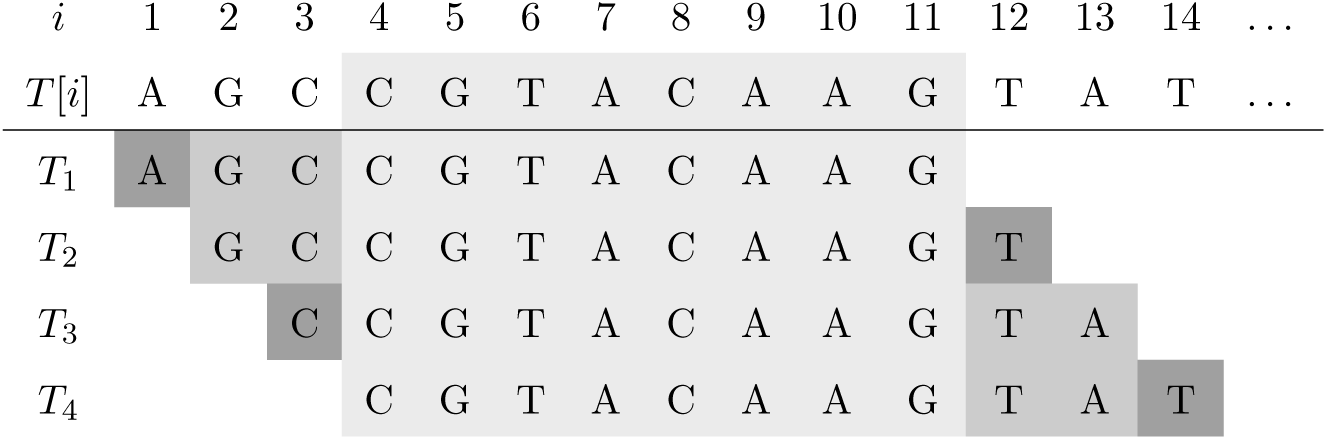
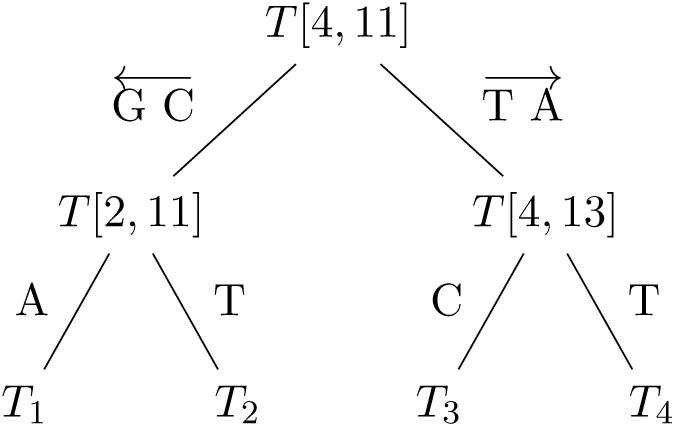
Searching *s* overlapping *k*-mers using optimum search schemes for the infix and extending it using back-tracking. Illustrated for *k* = 11 and *s* = 4. (a) First, the common overlap (light gray) is searched using optimum search schemes. Second, the search of *T*_1_ and *T*_2_ is continued recursively by extending the previously identified approximate matches of the infix in the index by GC to the left (allowing for the remaining number of errors; medium gray). *T*_1_ and *T*_2_ are then retrieved separately by backtracking in the index by one character to the left and one character to the right (allowing for an error, if any left; dark gray). *T*_3_ and *T*_4_ are extended analogously in a recursive manner. (b) The same strategy presented as a backtracking tree. It is traversed for all occurrences reported by the search of the infix *T* [4, 11] using optimum search schemes. Each edge also has to account for remaining errors, i.e., approximate string matching is performed using backtracking.

To further reduce the number of redundant computations, the set of overlapping *k*-mers is recursively divided into two sets of *k*-mers of roughly equal size that each share a larger common overlap among each other. This overlap is then searched using backtracking before the next recursive partitioning of *k*-mers. The recursion ends when a single *k*-mer is left and the number of occurrences can be reported and summed up, or no hits are found. The recursive extension is shown in figure 2b. Note, that there are two recursions involved: subdividing the set of *k*-mers and backtracking in each recursion step. Hence, the same partitioning steps and backtracking steps have to be performed for each set of preliminary matches represented by suffix array ranges in the FM index.

### 2.2 Approximate String Matching using Optimum Search Schemes

Backtracking performed in an index to search for approximate matches leads to an exponential running time in the number of errors. Especially allowing for errors at the beginning of the *k*-mer, i.e., branching at the topmost nodes in the backtracking tree is expensive. Hence, we use optimum search schemes [6] when searching the infix, a sophisticated search strategy that reduces the number of search steps performed in the index while still searching for all possible approximate matches.

Optimum search schemes are based on a framework by Kucherov et al. called search schemes that allows formalizing search strategies in a bidirectional index [7]. The sequence to be searched is split into *p* pieces and searched by certain combinations of the pieces in the index while trying to reduce the number of search steps performed in the index.

Formally, a search is a triplet *S* = (*π, L, U*) of integer strings each of length *p. π* is a permutation of the numbers {1, 2, *…, p*} indicating the order in which the pieces are searched. Starting from an arbitrary piece *π*[0] the subsequent pieces need to be adjacent to the previously searched pieces. *L* and *U* are non-decreasing integer strings indicating the lower and upper bound of errors. After the piece *π*[*i*] is searched a total number of *L*[*i*] and *U* [*i*] errors must have been spent. A set of searches that covers all possible error distributions with *e* errors and *p* pieces forms a search scheme. Intuitively speaking, approximate string matching is sped up by reducing the number of errors allowed in the first pieces of each search. By performing multiple searches starting with different pieces, it is guaranteed that all possible error distributions among the pieces are still covered.

Optimum search schemes are search schemes that are optimal under certain constraints, i.e., the number of backtracking steps in an index over all searches are minimized while still covering all error distributions. Figure 3 illustrates the optimum search scheme for *e* = 2 errors, *p* = *e* + 2 pieces and up to 3 searches.

**Figure 3:**
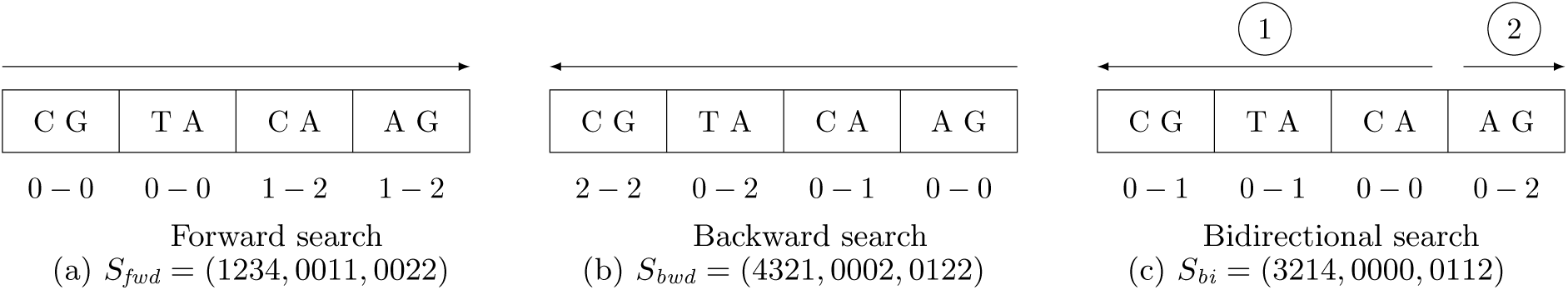
The optimum search scheme for 2 mismatches consists of 3 searches with 4 pieces each. The arrows indicate in which order the pieces are searched. The error bounds below each part are cumulative bounds, i.e. the minimum respectively maximum number of errors that *must* respectively *can* be spent until searching the end of the corresponding piece. Illustrated for searching the 8-mer CGTACAAG. The forward searches covers the error distributions 0010, 0011, 0020, the backward search covers 2000, 1100, 0200, 1010, 0110, and the bidirectional search 0000, 0001, 0002, 1000, 1001, 0100, 0101.

To allow searches that start with a middle piece as illustrated in figure 3c a bidirectional index is required again. To improve the overall running time of the index-based search we use a fast implementation of bidirectional FM indices based on EPR dictionaries [8].

The question remains on how to combine the improvements of sections 2.1 and 2.2, i.e., how to choose *s*, the number of adjacent *k*-mers that are searched together starting with their common sequence using optimum search schemes. On one hand approximate string matching using optimum search schemes is more efficient than simple backtracking, hence a longer common infix is favorable. On the other hand a longer common infix means fewer adjacent *k*-mers are searched at once which leads to more redundant search steps due to the high similarity of overlapping *k*-mers. GenMap chooses *s* according to the following equation derived from optimal values that were determined experimentally on different genomes such as the human and barley genome (see [9] for details). clamp(*v, l, r*) returns *v* if it lies within the range, i.e., *l ≤ v ≤ r*, and returns *l* or *r* if it is less or greater.

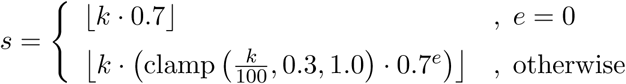

### 2.3 Skipping redundant k-mers

Finally, we avoid searching the same *k*-mer multiple times. Especially *k*-mers from repeat regions may occur many times without errors in the text. Since they all share the same frequency value, it should be avoided to compute it more than once. Hence, after searching and counting the occurrences of a *k*-mer, we locate the positions of the exact matches and set all their frequency values in *F* accordingly.

We observed that this strategy leads to longer runs of frequency values forwarded to positions with uncomputed frequency values. When we encounter forwarded frequency values of previously counted *k*-mers during the scan over the text, they can be skipped.

## 3. Benchmarks

At first we compare the running times for computing the frequency on the human genome for different lengths and errors based on Hamming distance. We ran GEM in its exact mode as well as with its approximation. For the latter the authors recommend *t* = 7. Table 1a compares the running times for shorter *k* that are of interest for applications such as identifying marker genes, presented in section 4. Table 1b shows typical instances used for applications in read mapping based on a typical Illumina read length. Even though longer Illumina read lengths are more common these days, we choose a shorter read length, since the frequency is easier to compute for longer *k*-mers and a fixed number of errors.

**Table 1:**
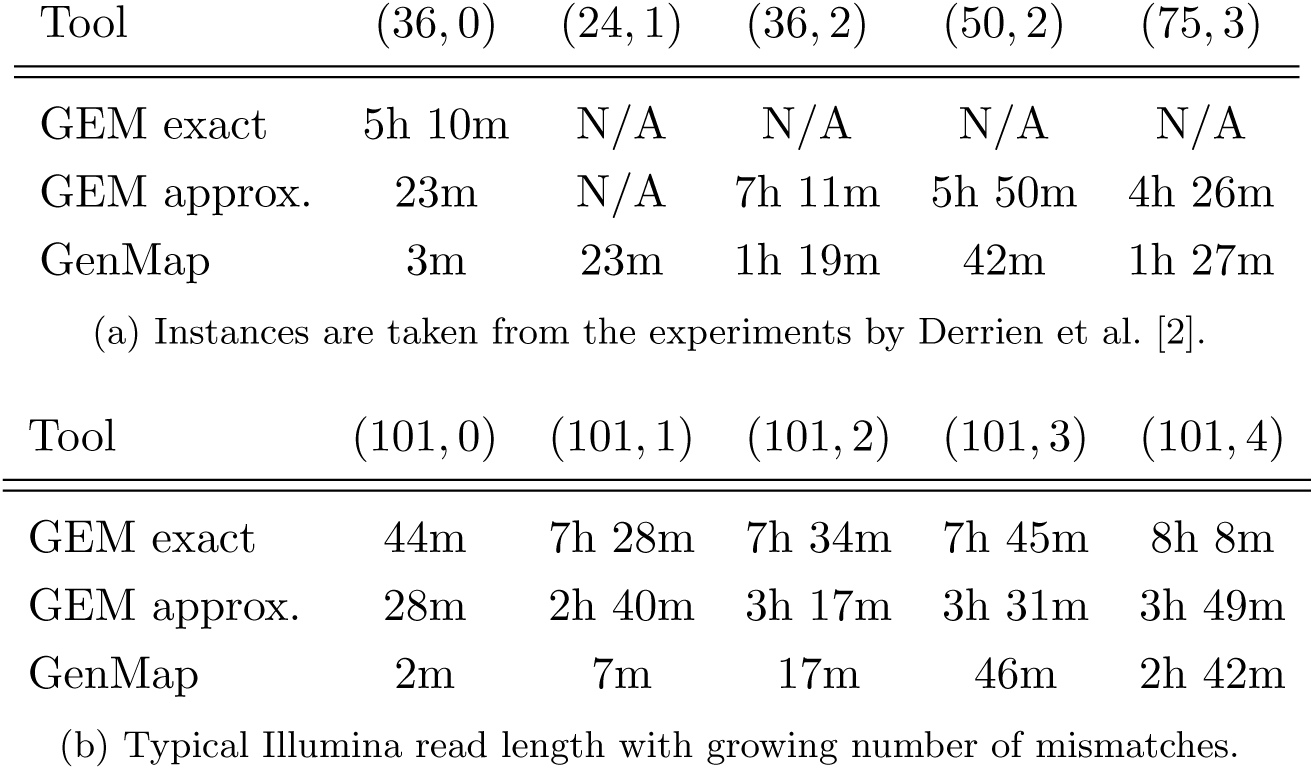
Running times for computing the frequency of the human genome (GRCh38) using 16 threads. Timeouts of 1 day are represented as N/A.

For all computed instances, GenMap is faster than GEM. Compared to the approximate mode we are almost a magnitude faster for smaller number of errors, but for 4 errors the heuristic of GEM pays off and is almost as fast as our algorithm. Interestingly, the increase of the running time of GEM in its exact mode gets smaller with more errors. For 101-mers with 1 to 4 errors the running time is always about 7 to 8 hours, nonetheless GenMap is still faster by a factor from 3 of up to 64 (4 and 1 errors). Even when searching without errors where no backtracking has to be performed, our tool is faster by a factor of 20 to 100 (for 101-mers and 36-mers). The most noticeable improvement is achieved for short *k*-mers. Derrien et al. point out that their algorithm is not suitable for small *k* and completely unfeasible for *k* < 30 without its approximation which is reflected by our benchmarks, whereas GenMap can handle these instances easily. GEM takes significantly longer, often does not even terminate within 24 hours on 16 threads.

GenMap is also faster than GEM when computing the frequency of small genomes like D. melanogaster. Since smaller genomes are generally less challenging, we omit the benchmarks here. For the human genome the memory consumption of GenMap is about 9 GB (using a bidirectional FM index with EPR dictionaries and a suffix array sampling rate of 10), while GEM takes up 4.5 GB (using an unspecified FM index implementation with a suffix array sampling rate of 32).

GenMap is also suitable to compute the frequency of larger and more repetitive genomes than the human genome. We computed the (50, 2)-frequency of the barley genome [10] as it contains large amounts of repetitive DNA [11]. Barley has 4.8 billion base pairs while the human genome has 3.2 billion base pairs. As expected the human genome has considerably more unique regions than the barley genome. To be precise 75.4 % of the 50-mers are unique in the human genome, and only 26.4 % in the barley genome. There are 12.0 % (54.4 %), 7.6 % (42.1 %) and 4.8 % (25.6 %) 50-mers in human DNA (resp. barley DNA) with at least 10, 100 and 1,000 occurrences. Computing the (50, 2)-frequency of barley on 16 threads took less than 1h 15m with GenMap and nearly a day with GEM using its heuristic with *t* = 6 (automatically chosen by GEM).

In conclusion, GenMap is a magnitude faster than GEM in its exact mode, and still faster than GEM using its heuristic, while GenMap is always exact. Even for up to 4 errors GenMap achieves a reasonable running time. This is due to the three techniques described in the previous section. Further improvements can be implemented which might speed up the algorithm even further, such as in-text verification [9], i.e., locating at some point the partially searched *k*-mers and verifying whether their locations in the text match the *k*-mer with respect to the error bound. A location and verification step in the text is often several times faster than finishing the index-based search.

All tests were conducted on Debian GNU/Linux 7.1 with an Intel Xeon E5-2667v2 CPU. To avoid dynamic overclocking effects in the benchmark the CPU frequency was fixed to 3.3 GHz. The data was stored on a virtual file system in main memory to avoid loading it from disk during the benchmark which might affect the results due to I/O operations.

We used the only available version 1.759 beta of the GEM suite that included the mappability program. We did not reach the authors for other versions including the mappability tool. Other available and newer versions do not offer this feature anymore. The running times we measured for GEM approx. differs considerably from the running times for GRCh37 published by the authors. Even when we ran it on a similar CPU with the same number of cores we were 2 to 5 times slower than their published benchmarks. One reason might be that the only available version of GEM with the mappability functionality was published as a beta version, however it was a year after their paper. Nonetheless, GenMap is still faster than the running times published by Derrien et al. For a fair comparison in our benchmark we reduced the genomes to the dna4 alphabet, i.e., replaced Ns by random bases. Based on tests we observed that GEM neither computes the mappability of *k*-mers that have unknown bases, nor considers them as mismatches in its default mode even when errors are allowed.

A more recent tool to compute the mappability is Umap [12]. It is limited to computing the (*k,* 0)-mappability and reporting only unique *k*-mers, i.e., regions with a mappability value of 1. It uses the read mapper Bowtie to search every single *k*-mer in the genome and filter non-unique *k*-mers afterwards. Due to the constraints we excluded it from our benchmarks. From the authors benchmarks we can conclude that GenMap still outperforms Umap as GenMap needs less than one hour without parallelization to compute the (*k,* 0)-mappability (see table 1), while Umap needs about 200 hours.

## 4. Experiments

Although the main focus of this work lies on presenting a new and fast algorithm for computing the mappability of a genome, we propose an application for identify marker genes illustrated by a small example on E. coli strains. Marker genes or marker sequences are short subsequences of genomes whose presence or absence allow determining the organism, species or even strain when sequencing an unknown sample or help building phylogenetic trees [13]. Depending on the marker length it can span up to dozens of reads. Instead of assembling the strain to search for marker genes or applying experimental methods such as PCR-based AFLP (amplified fragment length polymorphism) [14], we propose using its mappability. GenMap can be used to compute the mappability on a set of sequences, i.e., each *k*-mer is searched and counted in all strains at once. We consider two use cases: on the one hand we want to identify *k*-mers that match a sequence uniquely to determine the exact strain. On the other hand, we want to search for *k*-mers shared by many or all strains in the same phylogenetic group.

Small adjustments to the mappability algorithm are necessary. The frequency vector can be computed on multiple strains at once, but the algorithm does not distinguish whether a *k*-mer of some strain matches the same strain multiple times or different strains. While the original algorithm simply counts the occurrences, we are now interested in the number of different strains it matches for finding marker sequences. A *k*-mer with a frequency greater than 1 can still be a suitable marker sequence if it only matches one of the strains. We also extended GenMap to output a CSV file with the locations of each *k*-mer which can be parsed to filter *k*-mers that only occur in a subset of strains.

To test this approach we used a data set of E. coli strains. It was shown that E. coli can be grouped into four major phylogenetic groups (A, B1, B2, and D) [15]. The authors identified two marker genes (chuA and yjaA) and an anonymous DNA fragment (TspE4.C2) whose combination of presence or absence in the genome can determine the phylogenetic group.

We computed the (30, 2)-mappability on four different strains of group B1^1^. According to the study all strains within B1 share the anonymous DNA fragment TspE4.C2 of 152 base pairs. We used GenMap to search for both, unique *k*-mers among all strains as well as *k*-mers that occur in each strain at least once, see figure 4a for an illustration. We observed that TspE4.C2 is an exact match in all strains and the 30-mers in this region also have a mappability value of exactly 0.25 when accounting for 2 errors. We further found numerous 30-mers with a mappability of 1, thus allowing to determine a strain among those four, while still accounting for sequencing errors and mutations. Table 2 lists the number of *k*-mers identified. We counted the number of *k*-mers matching only one strain, i.e., the strain the *k*-mer originated from. We refer to this count as *unique*. Additionally, we counted how many of these *k*-mers matched multiple times to the strain, referred to as *pseudo*. GenMap allows to exclude these pseudo marker genes when computing the mappability on multiple sequence files, i.e., it is only counted in how many sequence files a *k*-mer is present. To avoid counting highly overlapping *k*-mers in large unique regions, we break down the numbers for non-adjacent *k*-mers as well, i.e., for a *k*-mer to be considered it must have a preceding *k*-mer with a mappability value smaller than 1.

**Table 2:** (30, 2)-mappability on four strains of E. coli assigned to the phylogenetic group B1 based on the known marker genes by Clermont et al. We computed the mean distance of the unique marker sequences and their standard deviation.

| Strain | all $k$-mers | | | non-adjacent $k$-mers | | |
| --- | --- | --- | --- | --- | --- | --- |
| | Unique | Pseudo | ∅ Dist. | Unique | Pseudo | ∅ Dist. |
| IAI1 | 171,942 | 4,992 | $27 \pm 627$ | 1,829 | 81 | $2,476 \pm 5,560$ |
| SE11 | 305,439 | 10,365 | $15 \pm 447$ | 2,356 | 176 | $1,942 \pm 4,708$ |
| 11128 | 260,305 | 40,101 | $20 \pm 953$ | 2,494 | 685 | $2,049 \pm 9,517$ |
| 11368 | 434,033 | 108,968 | $13 \pm 912$ | 3,142 | 1,116 | $1,674 \pm 10,592$ |

**Figure 4:**
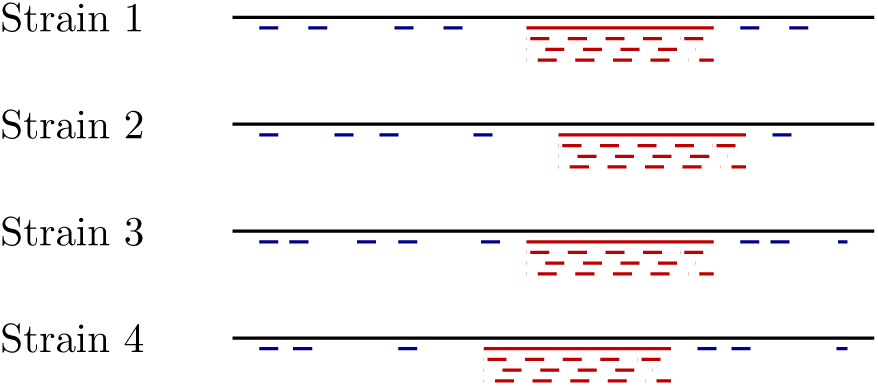
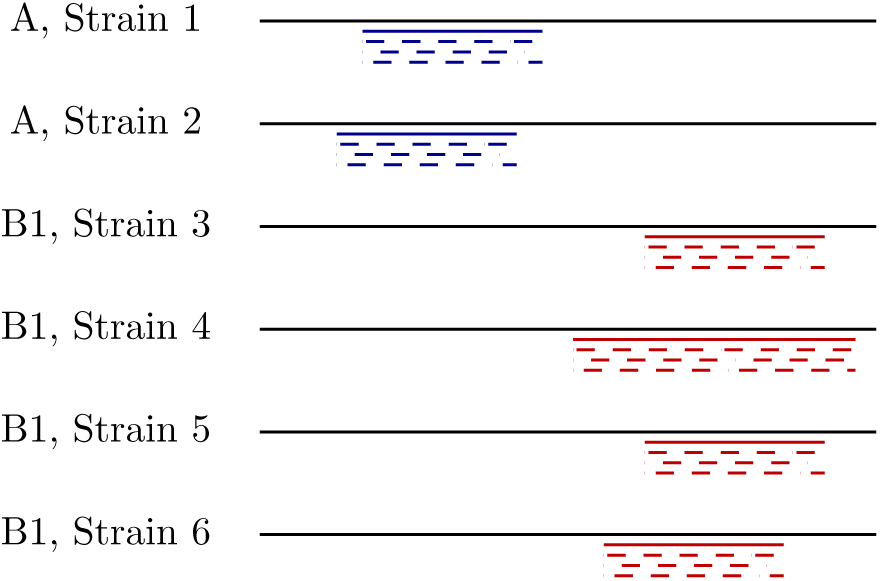
Illustration of the experiments performed on E. coli sequences in tables 2 and 3. (a) Four strains belonging to the same phylogenetic group. The sequence in red is conserved within this group and a marker gene. The red *k*-mers belonging to this marker gene are also all found in the other strains. The *k*-mers in blue are unique among all four strains and allow distinguishing each of the strains. (b) Six sequences belonging to two different phylogenetic groups. Marker sequences are highlighted in red and blue. They only occur in one of the groups and are present in all of its strains.

In table 3 we present the data of a second experiment, where we select strains from more than one group (A and B1), see figure 4b for an illustration. Again, we computed the (30, 2)-mappability, but this time we counted *k*-mers that match all strains in one group but no strain in the other group.

**Table 3:**
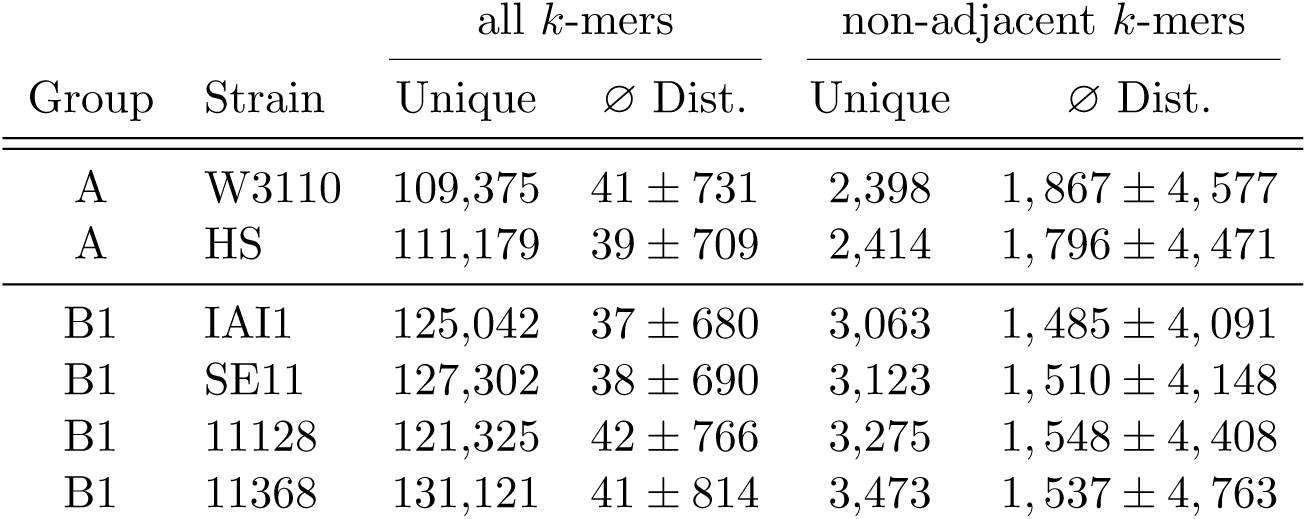
(30, 2)-mappability on six strains of E. coli of the groups A and B1. Only *k*-mers were counted that perfectly separated the strains in A from B1, i.e., if and only if the *k*-mer matched all strains of A and no strain of B1 and vice versa.

| Group | Strain | all $k$-mers | | non-adjacent $k$-mers | |
| --- | --- | --- | --- | --- | --- |
| | | Unique | ∅ Dist. | Unique | ∅ Dist. |
| A | W3110 | 109,375 | $41 \pm 731$ | 2,398 | $1,867 \pm 4,577$ |
| A | HS | 111,179 | $39 \pm 709$ | 2,414 | $1,796 \pm 4,471$ |
| B1 | IAI1 | 125,042 | $37 \pm 680$ | 3,063 | $1,485 \pm 4,091$ |
| B1 | SE11 | 127,302 | $38 \pm 690$ | 3,123 | $1,510 \pm 4,148$ |
| B1 | 11128 | 121,325 | $42 \pm 766$ | 3,275 | $1,548 \pm 4,408$ |
| B1 | 11368 | 131,121 | $41 \pm 814$ | 3,473 | $1,537 \pm 4,763$ |

This example shows that mappability on multiple species or strains can be used to identify possible marker sequences. Short *k*-mers could be used to search a data set of reads instead of searching for marker genes that span multiple reads. Since computing the (30, 2)-mappability on a few E. coli strains even takes less than a minute on a consumer laptop, this method is suitable to be run on large sets of similar E. coli strains to identify new marker sequences, even with errors accounting for uncertainty arising from sequencing and mutations such as SNPs.

## 5. Discussion and Outlook

We have presented GenMap, a fast and exact algorithm to compute the mappability of genomes up to *e* errors, which is based on the C++ sequence analysis library SeqAn [16]. It is significantly faster, often by a magnitude than the algorithm from the widely used GEM suite while refraining from approximations.

Mappability has already been used for various purposes [2]. In this paper we proposed a new application, the computation of mappability on a set of genomes to identify marker genes for grouping and distinguishing genomes by short *k*-mers and illustrated it with a small example on closely related E. coli strains.

The ability to compute the exact mappability opens up new applications such as incorporating the mappability information during read mapping instead of the post-processing phase. In [9] we show that new read mapping strategies can lead to faster mapping. During the index-based search of a read the possible locations of the eventually completely mapped read can be examined beforehand to filter repetitive regions without repeat masking. This allows for new mapping strategies to improve the running time of state-of-the-art read mappers and reduce post-processing overhead.

## Acknowledgements

The authors acknowledge the support of the de.NBI network for bioinformatics infrastructure, the Intel SeqAn IPCC and the IMPRS for Computational Biology and Scientific Computing.

## Footnotes

[1 Strains and GenBank accession numbers of their assemblies: IAI1 O8 (GCA 000026265.1), SE11 O152:H28 (GCA 000010385.1), 11128 O111:H-(GCA 000010765.1), 11368 O26:H11 (GCA 000091005.1)

